# Zika Fetal Neuropathogenesis: Etiology of a Viral Syndrome

**DOI:** 10.1101/050674

**Authors:** Zachary A. Klase, Svetlana Khakhina, Adriano De Bernardi Schneider, Michael V Callahan, Jill Glasspool-Malone, Robert Malone

**Affiliations:** Department of Biological Sciences, University of the Sciences, Philadelphia, PA, USA; Department of Bioinformatics and Genomics, University of North Carolina at Charlotte, Charlotte, NC, USA; Department of Medicine, Division of Infectious Diseases, Massachusetts General Hospital, Boston, Massachusetts, USA; Atheric Pharmaceutical, Scottsville, VA, USA; Class of 2016, Harvard Medical School Global Clinical Scholars Research Training Program, Boston, Massachusetts, USA

## Abstract

The ongoing Zika Virus epidemic in the Americas, and the observed association with both fetal abnormalities (primary microcephaly) and adult autoimmune pathology (Guillain-Barré syndrome) has brought attention to this neglected pathogen. While initial case studies generated significant interest in the Zika virus outbreak, larger prospective epidemiology and basic virology studies examining the mechanisms of Zika viral infection and associated pathophysiology are only now starting to be published. In this review, we analyze Zika fetal neuropathogenesis from a comparative pathology perspective, using the historic metaphor of “TORCH” viral pathogenesis to provide context. By drawing parallels to other viral infections of the fetus, we identify common themes and mechanisms that may illuminate the observed pathology. The existing data on the susceptibility of various cells to both Zika and other flavivirus infections are summarized. Finally, we highlight relevant aspects of the known molecular mechanisms of flavivirus replication.

**Key Learning Points:**

1. Viral TORCH pathogens reveal common patterns of fetal pathophysiology and vertical transmission which are relevant to Zika Virus fetal neuropathogenesis.
2. The teratogenic effects of Zika Virus infection during the first trimester may involve infection of the trophoblast, viral translocation across the placenta, migration of infected cells resulting in embryonic infection, or indirect effects associated with high levels of inflammatory cytokines produced by infected placenta.
3. Pre-existing maternal non-neutralizing antibody to Zika virus may enhance the probability of infection or more severe disease in the fetus.
4. AXL has been identified as a major receptor for Zika Virus.
5. Zika virus activation of Toll Like Receptor 3 (TLR-3) pathways in central nervous system cells may trigger apoptosis and attenuate neurogenesis, directly contributing to fetal neuropathology.
6. Flaviviruses subvert host autophagy and noncoding RNA regulatory pathways.
7. Recognition of viral sequences by regulatory RNA binding proteins such as Musashi may have a role in Zika pathogenesis and host tissue tropism.
8. Evidence from other TORCH viral pathogen studies indicate multiple plausible hypotheses for transplacental infection by Zika virus during the second or third trimester, including transcytosis of non-neutralizing antibody-coated Zika virus complexes.

**Key References:**

Adibi JJ, Marques ET Jr, Cartus A, Beigi RH. Teratogenic effects of the Zika virus and the role of the placenta. Lancet 2016; 387: 1587–90 (Hypothesis)
Adams Waldorf KM, McAdams RM. Influence of infection during pregnancy on fetal development. Reproduction. 2013 Oct 1;146(5) (Review)
Hamel R, Dejarnac O, Wichit S, Ekchariyawat P, Neyret A, Luplertlop N, et al. Biology of Zika Virus Infection in Human Skin Cells. J Virol. 2015;89(17):8880–96.
Mlakar J, Korva M, Tul N, Popović M, Poljšak-Prijatelj M, Mraz J, et al. Zika Virus Associated with Microcephaly. N Engl J Med. 2016 Feb 10.
Paul LM, Carlin ER, Jenkins MM, Tan AL, Barcellona CM, Nicholson CO, Trautmann L, Michael SF, Isern S. Dengue Virus Antibodies Enhance Zika Virus Infection. bioRxiv doi: http://dx.doi.org/10.1101/050112
Crow YJ, Manel N. Aicardi-Goutieres syndrome and the type I interferonopathies. Nat Rev Immunol. 2015;15(7):429-40.
Tonduti D, Orcesi S, Jenkinson EM, Dorboz I, Renaldo F, Panteghini C, et al. Clinical, radiological and possible pathological overlap of cystic leukoencephalopathy without megalencephaly and Aicardi-Goutieres syndrome. Eur J Paediatr Neurol. 2016.
Cipolat Mis MS, Brajkovic S, Frattini E, Di Fonzo A, Corti S. Autophagy in motor neuron disease: Key pathogenetic mechanisms and therapeutic targets. Molecular and Cellular Neurosciences. 2016;72:84-90.
Dang J, Tiwari SK, Lichinchi G, Qin Y, Patil VS, Eroshkin AM, Rana TM. Zika Virus Depletes Neural Progenitors in Human Cerebral Organoids through Activation of the Innate Immune Receptor TLR3. Cell Stem Cell. 2016: 19: 1–8.
Vianna FS, Schuler-Faccini L, Leite JC, de Sousa SH, da Costa LM, Dias MF, et al. Recognition of the phenotype of thalidomide embryopathy in countries endemic for leprosy: new cases and review of the main dysmorphological findings. Clin Dysmorphol. 2013;22(2):59-63.

---

Zika Virus (ZIKV), a mosquito vectored flavivirus, was first isolated in 1947 from a sentinel research monkey caged in the Zika forest canopy within Uganda (1, 2). Soon after discovery, ZIKV was observed to infect humans (3). Travel, shipping, and the worldwide distribution of human hosts and mosquito vectors (including *Aedes aegypti* and other *Aedes* species) has facilitated a global radiation of Zika viral infection (4). More recently, introduction of ZIKV into naïve human populations has yielded rapidly spreading outbreaks in various Pacific island clusters (Cook Island, Easter Island, French Polynesia and Micronesia) and the ongoing epidemic in the Americas which may have originated in Haiti (5), and has subsequently spread throughout Brazil, the Caribbean, and worldwide via travelers visiting affected regions (6, 7). In ZIKV endemic regions such as continental Africa and Asia, there is epidemiologic support for the hypothesis that people are exposed to ZIKV during childhood and thereby develop immunity prior to puberty in both males and females. Introduction of ZIKV into dense immunologically naïve populations has facilitated rapid viral evolution, including conserved modifications consistent with possible adaptation to a human host (8, 9). Most pertinent to the current concern about ZIKV is the infection of pregnant women who are immunologically naïve to ZIKV, intrauterine infection of their babies, and associated increased risk of congenital malformations consistent with other fetal pathogens such as those historically referred to by the TORCH acronym (**T**oxoplasmosis, **O**ther (HIV, Syphilis, Varicella Zoster Virus (VZV) etc.), **R**ubella, **C**ytomegalovirus (CMV) and **H**erpes simplex virus-2 (HSV)).

ZIKV fetal syndrome resembles but is more severe than that observed with many other intrauterine viral infections. Typical presentation includes multiple defects; microcephaly, facial disproportionality, cutis gyrata, hypertonia/spasticity, hyperreflexia, and irritability; abnormal neurologic image findings include coarse and anarchic calcifications mainly involving the subcortical cortical transition and the basal ganglia, ventriculomegaly secondary to the lack of brain tissue, and lissencephaly (7, 10-13). This alarming and consistent clinical presentation provoked a rapid regional mobilization of public health experts in Pernambuco (in the Northeast of Brazil). Investigation soon revealed a correlation between ZIKV infection and the unusually high rate of infant microcephaly observed at the heart of the outbreak in Recife, Pernambuco. The striking features of ZIKV fetal syndrome may have gone unrecognized during prior outbreaks in the Pacific islands, or may involve regional confounding variables or risk cofactors present in Brazil such as prior exposure to Dengue virus (14, 15). The current pathology may also be consequent to recent viral mutations, such as observed changes in the prM protein of the Brazilian ZIKV strains (8, 16, 17). It has been demonstrated that ZIKV can infect human induced pluripotent stem cell -derived neural progenitor cells as well as human neurospheres and brain organoids in vitro, resulting in dysregulation of cell-cycle-related pathways and increased cell death (18-21). While the etiology remains unconfirmed, there appears to be a shift in the spectrum and incidence of birth defects between the latter stage of the French Polynesian outbreak (22) and what is now being observed in Recife, Rio, and throughout northern Brazil and surrounding regions (23, 24). In general, the combination of epidemiologic association and experimental research results strongly support a causal relationship between intrauterine ZIKV infection and fetal primary microcephaly.

Historically, human infection with ZIKV has presented in adults and young children as a mild, self-limiting, non-life threatening infection with clinical symptoms appearing in 20% of infected patients, and up to 80% being clinically asymptomatic during initial infection. Symptoms typically persist an average of 4 to 5 days to approximately one week from initial onset of headache and fever. Key major symptoms following retro-orbital and frontal headache and fever include a less consistent presentations of malaise, arthalgias, conjunctivitis, and pruritic maculopapular rash. More severe causes include escalation of the symptoms above, as well as nausea, vomiting and GI distress (4). The most recent assessment of clinical signs and symptoms of acute Zika virus infection observed in Puerto Rico includes rash (74%), myalgia (68%), headache (63%), fever (63%), arthralgia (63%), eye pain (51%), chills (50%), sore threat (34%), petechiae (31%), conjunctivitis (20%), nausea/vomiting (18%), and diarrhea (17%) (25). Based on blood bank screens, viremia can begin up to 10 days before onset of symptoms (26), and the modest plasma viral titers observed often clear within two days of presentation with clinical symptoms, similar to what is observed with Dengue (27). At present, definitive diagnosis requires a polymerase chain reaction (PCR)-based test, and development of a rapid serologic diagnostic test is complicated by antibody cross-reactivity with other co-circulating arboviruses (28, 29). Historic serologic surveillance studies have been compromised by acute Zika infection induction of high titers of anti-dengue and even anti-chikungunya convalescent IgG levels, routinely at titers above 1:1280 (30, 31).

Current best estimates for the basic reproductive ratio (R_0_) for ZIKV varies between 1.2 and 6.6 (32-34), with seroconversion rate being approximately 70%, upon achieving maximal herd immunity. This limitation on further infection within a naïve population is typically achieved within four to eighteen months of initial introduction (35, 36). Acute motor axonal neuropathy-type Guillain-Barré syndrome (GBS) occurred at a rate of 1 in 5,000 cases of ZIKV during the outbreak in French Polynesia (15); the rate for GBS and all combined neurologic disease in the Americas may be as high as 1 in 100 cases (25). A clear temporal relationship between the peak of Zika virus infection in a susceptible population and a peak of GBS incidence following five to nine weeks later has been demonstrated, consistent with an autoimmune-mediated (rather than direct viral infectious neuropathy) pathologic mechanism (37). Interim analysis of an ongoing prospective case study of ZIKV -infected pregnancies indicates a birth defect rate of circa 29% (23). For the sake of illustration, the potential impact of these epidemiologic estimates on the anticipated 2017-2018 Puerto Rico birth cohort is summarized in Figure 1.

**Figure 1.**
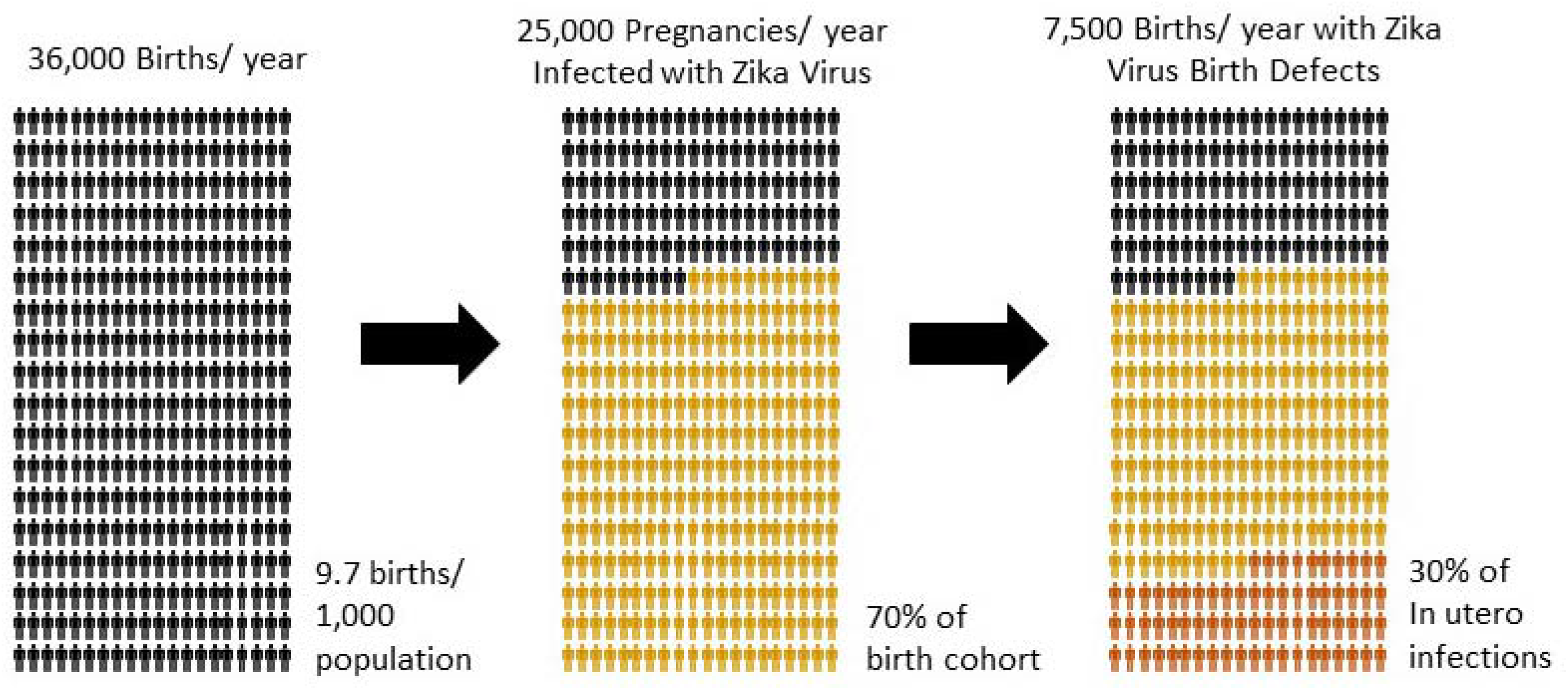
**Projected teratogenic impact of maternal ZIKV infection on 2017-2018 birth cohort, Puerto Rico**. For illustration purposes, the potential impact of unencumbered ZIKV spread through Puerto Rico on the cumulative one-year incidence of ZIKV-associated birth defects has been estimated and graphically summarized. Birth defect rate is based on preliminary data involving defects visible by *in utero* ultrasound examination from Brazilian (Rio) prospective pregnancy cohort study (23). Final seroconversion rate of 70% is based on seroconversion observed with prior island outbreaks in Yap and French Polynesia (35, 36). Annual birth cohort for Puerto Rico is approximated as 36,000 infants, a number which presumes that the incidence of pregnancy is not impacted by anticipated risk of ZIKV infection or public health policy recommendations. Total birth defect rate associated with intrauterine ZIKV infection in Northern and Central Brazil is currently not determined, and may exceed 30% of all Zika-infected pregnancies.

In the current outbreak in the Americas, there is evidence for sexual transmission of the virus (38-41). While ZIKV RNA can be detected in breast milk, urine, semen and sputum from infected individuals (42), replication competent virus has been most readily cultured from semen samples. Semen ZIKV RNA levels may be up to 100,000 times higher than corresponding plasma levels (43). Preferential ZIKV replication in testes has been hypothesized. ZIKV is shed in semen for an extended period, and the average duration of shedding has yet to be determined (43). The stability of ZIKV in aqueous suspension, on surfaces or as fomites is unknown, but other flaviviruses can persist under various ambient conditions for extended periods (44-48). Zika virus sequences have been difficult to detect in trapped mosquitoes from outbreak areas, but have recently been recovered from *Aedes albopictus* mosquitoes by the Laboratory of the Institute of Epidemiological Diagnosis and Reference (InDRE), which functions as part of the Mexico Epidemiological Surveillance System (SINAVE) (49). ZIKV is more stable than Dengue virus (16), and so it cannot be assumed that sexual transmission is the only means of direct human to human infection. Sequence comparisons of ZIKV isolates indicate significant genetic differences between historic samples obtained from mosquito species and more modern isolates from human sources, including human samples obtained during the current outbreak in the Americas (8, 9). Any clinical significance associated with these viral genetic changes has yet to be elucidated.

The apparent teratogenic effects of ZIKV infection have turned what was once considered a relatively benign pathogen into a subject of great social and scientific concern. Detection of ZIKV RNA and particles in amniotic fluid and fetal brain obtained from the products of conception strongly suggest that the virus is capable of directly infecting fetal tissue (12, 13). When considering the vast array of human pathogens, the probability of a mother passing an infection to her developing fetus is relatively rare. However, examples of pathogens consistently capable of vertical intrauterine transmission do exist, and can be associated with teratogenic effects. These viral diseases involving intrauterine infection may illuminate and inform research into the possible mechanisms by which ZIKV may induce fetal neuropathology as well as other birth defects, and may facilitate development of public health risk mitigation strategies and potential treatments.

## TORCH Viral Pathogens

Teratogenic infectious agents that are vertically transmitted from mother to infant during pregnancy, childbirth or breastfeeding have traditionally been classified as TORCH pathogens. For the purpose of this review we will focus on the classical viral TORCH pathogens: Rubella, CMV, HSV and VZV. These viruses can cross the placenta and cause congenital defects including, but not limited to, microcephaly, growth and mental retardation, heart disease, hearing loss and blindness (50-52). Years of scientific research concerning TORCH pathogen infection and teratogenicity have yet to identify therapeutic interventions which reduce occurrence of serious medical sequela and miscarriages for most of these viruses. Current preventative measures are limited to vaccination and avoiding viral exposure, or dosing with acyclovir for HSV (53). These approaches have limitations, and are not globally available. The most extensive fetal damage associated with viral TORCH infections typically takes place when the mother is infected during first eight weeks of the pregnancy, during which time the central nervous system (CNS) of the developing fetus is actively forming. With most viral TORCH pathogens, birth defect risk and severity is significantly reduced when infection occurs after seventeen weeks of gestation (54). Often first trimester infections result in miscarriages. Not all fetal congenital abnormalities manifest clinically at birth, and may present later in a child’s development. As summarized in Table 1, presence of congenital defects at birth is typically linked to TORCH infection at earlier stages of gestation.

**Table 1.** **Selected Viral TORCH pathogens and associated morbidity**. After (54).

| Viral TORCH Pathogen | Symptoms | First or Second Trimester Teratogen | Third Trimester Teratogen | Primary microcephaly | Spontaneous abortion or fetal death |
| --- | --- | --- | --- | --- | --- |
| **Rubella virus**<br>**(German measles)** | Defects in multiple organ systems including the ophthalmic (cataracts and microphthalmia), cardiac, neurological (deafness, mental retardation), and increased risk of type 1 diabetes in childhood | + | - | + | + |
| **Cytomegalovirus** | Mental retardation, sensorineural hearing loss, jaundice, hepatosplenomegaly, petechiae, preterm birth, preeclampsia, and fetal growth restriction | + | - | + | + |
| **Herpes simplex virus** | Encephalitis, sepsis, cataracts, pneumonitis, myocarditis, hepatosplenomegaly, chorioretinitis, encephalitis, and mental retardation | + | + | + | + |
| **Varicella zoster virus**<br>**(chickenpox)** | Skin lesions, neurological and eye defects, limb hypoplasia, fetal growth restriction, and defects of multiple organ systems | + | - | +/- | + |
| **Zika virus** | Microcephaly, facial disproportionality, cutis gyrata, hypertonia/spasticity, hyperreflexia, and irritability; abnormal neuroimages include calcifications, ventriculomegaly, and lissencephaly | + | + | + | + |

## Rubella (German measles)

Prior epidemic outbreaks of Rubella and consequent associated birth defects may provide the best illustration of the neonatal health risks of the current ZIKV outbreak in the Americas (55), although the incidence of congenital rubella syndrome (CRS) associated with initial outbreaks in Rubella naïve populations (56) appears to have been significantly less than what is being documented with ZIKV in Brazil (23). Rubella virus (RuV) is a member of the *Rubivirus* genus and *Togaviridae* family. The Rubella genome is encoded on a positive single stranded RNA (ssRNA), which is assembled on a protein scaffold and surrounded by lipid envelope. Host cell infection with RuV is driven by two glycoproteins, E1 and E2. Encoded in by the RuV genome, these glycoproteins assemble as heterodimers on the surface of the viral envelope and function similarly to the fusion proteins of flaviviruses (57, 58). E1 protein trimer directly inserts into host cell plasma membrane lipid bilayer, and using hairpin motion, brings the RuV closer to the cell surface to facilitate endocytosis (59, 60). The release of viral genome into the host cell occurs via low pH, and with the Ca^2+^ dependent E1 trimer conformational changes associated with maturing endocytic vesicles (61). Recent work has identified myelin oligodendrocyte glycoprotein as a receptor with affinity for RuV E1 protein (62). This discovery may provide a causal link between rubella virus and brain damage in fetuses with CRS. RuV infection of pregnant women has a pronounced teratogenic effect, especially during the first gestational trimester (59). Pathological and immunohistochemical analyses of aborted fetuses with CRS demonstrated wide spread necrosis to organs including eye, heart, brain and ear, and are associated with the presence of rubella virus in all tissues (63). In-vitro studies suggest that RuV infection inhibits normal growth and differentiation of human embryonic mesenchymal cells (64). RuV encoded replicase P90 protein has been shown to disrupt actin cytoskeleton formation by directly binding and inhibiting Cytron-K kinase, a cytokinesis regulatory protein (65). Inhibition of Cytron-K leads to cell cycle arrest and apoptosis in developing neuronal populations and retina of *in vitro* cultured mouse embryos (66). Additionally, Rubella virus infection of placenta and embryonic cells induces interferon expression, especially in the placenta (67). The most commonly observed outcomes of CRS are congenital cataracts (97.4%), inner ear abnormalities (73.9%), microcephaly (68.4%), and congenital heart defects (57.9%) (52, 63, 68, 69). If the infection occurs during the first trimester, the rate of CRS is 80-90%. Odds of intrauterine development of extensive CRS dramatically decreases after 12 weeks of gestation (50).

## Cytomegalovirus (CMV)

CMV is a member of the *Herpesviridae* family, *Betaherpesvirinae* subfamily and is also known as Human Herpesvirus 5 (HHV-5). Intrauterine CMV infection is linked to development of severe neurological handicaps, microcephaly (36%), intracranial calcifications, microgyria, eye defects and sensorineural hearing loss (68, 70-72). Congenital CMV infections are associated with radiographic findings which vary with gestational age at time of infection. Lissencephaly, including thin cerebral cortices, extremely diminished volume of white matter, delayed myelination, small cerebella, and very enlarged lateral ventricles have been correlated with CMV infection prior to eighteen weeks of gestational age, whereas those cases of congenital CMV infection which present with more normal gyral patterns (normal cerebral cortices, slightly diminished volume of white matter, delayed myelination, normal cerebella, and slightly enlarged lateral ventricles) are associated with third trimester infection (73, 74). These findings are similar to those observed with heritable disorders including cystic leukoencephalopathy without megalencephaly, Aicardi-Goutières syndrome, type 1 interferonopathies and RNASET2-related leukodystrophy (75, 76).

CMV is a double stranded DNA virus (dsDNA) with a complex envelope structure of 12 glycoproteins. Due to this complexity CMV, can bind to a broad spectrum of cell surface receptors, and quickly becomes ubiquitous in the human host after initial infection (77). CMV glycoprotein gB and heterodimer gM/gN have affinity to heparan sulfate proteoglycans (HSPGs), which are abundantly present on the surface of most cell types (78, 79). Additionally, CMV has been shown to bind epidermal growth factor receptor (EGFR) and β1 integrin coreceptors thereby facilitating proximity to the host cell membrane (80, 81).

CMV crosses host cell barriers via membrane fusion mediated by the gH/gL/gO and gB viral envelope glycoproteins (80, 82). CMV infection is mostly asymptomatic in immune competent adults, and forms a life-long latent infection. Primary CMV infection during pregnancy yields the highest risk of vertical transmission (32%) relative to virus re-activation in chronically infected mothers (1.4%) (83). CMV infection of the cytotrophoblast progenitor cells associated with floating villi in the placenta appears to elicit a shift in the Th1/Th2 cytokine balance of amniotic fluid and placental tissues, towards a Th1 profile, by upregulation of pro-inflammatory cytokines like MCP-1 and TNF-α (84, 85). This shift has been hypothesized to directly induce defects in placental formation and congenital abnormalities. There is significant evidence supporting the hypothesis that CMV virions transit placental barriers to fetal infection by co-opting the neonatal Fc receptor-mediated transport pathway for IgG (transcytosis) (86). However, replication of CMV in uterine endothelial cells may be required for subsequent infection of cytotrophoblasts (87, 88).

## HSV (HSV-1 and HSV-2)

HSV is a dsDNA enveloped virus belonging to the *Herpesviridae* family. Similar to CMV, HSV has a large number of glycoproteins present on the surface of its viral envelope, and can bind to multiple host cell receptors (89). HSV infection leads to formation of oral (HSV-1) and genital (HSV-2) lesions in adults. HSV host cell entry requires viral glycoprotein (primarily gD) binding to heparan sulfate and HveA (Herpes Virus Entry Mediator (HVEM) receptor), HveB (nectin-2) or HveC (nectin-1) receptors on the host cell plasma membrane surface. HSV enters the host-cell via membrane fusion or endocytosis (89). HSV can enter the CNS of adults, and in rare cases has been associated with clinical encephalitis (90). HSV infects neuronal cells through the nectine-1 receptor, and can form a latent and immunologically privileged reservoir of infection in the brain (91).

In contrast to CMV, cross-placental transition of HSV from mother to fetus is uncommon (92). Cells of the outer layer of the placenta do not express HveA, HveB or HveC, and cannot be infected by HSV (93). Congenital HSV infection is very rare, and usually occurs when a serologically negative mother is exposed to the virus during the first trimester of pregnancy. Congenital HSV pathology includes multi-organ failure, liver necrosis, encephalitis, microcephaly (32%), hydrocephalus, chorioretinitis and skin lesions (94, 95). HSV infection of placenta-associated cells induces inflammation and necrosis of placental tissue (94). Neonatal HSV-2 infection during childbirth or HSV-1 infection during the first year of life is more common, and is associated with up to 40% mortality. Aggressive anti-HSV treatment of neonates with acyclovir often controls the virus at the cost of long-lasting health risks to the child (96). There is a higher risk for HSV infection of the infant during childbirth in mothers that acquired genital HSV during the last trimester (^~^50%), while peripartum HSV-2 reactivation is associated with less than 1% of neonatal infections (96). This result suggests the role of maternal antibodies in protection of the child from HSV infection during birth. Congenital HSV infection is differentiated from perinatal infection by early onset (within 24h of birth) and increased severity of the symptoms (50). The relatively rare event of HSV microcephaly is exclusively associated with congenital infections (95).

## VZV (Chickenpox)

Varicella Zoster Virus (VZV) is a dsDNA enveloped virus. It belongs to *Herpesviridae* family, *Alphaherpesviridae* subfamily. VZV and HSV belong to the same subfamily, and share many characteristics (97). Similar to HSV, VZV can cause encephalitis, and can also form latent viral reservoirs in the brain (90, 98). The VSV viral envelope glycoprotein gE is essential for infection. This protein binds the Insulin-Degrading Enzyme (IDE) receptor, and employs heparan sulfate to facilitate host cell infection (99). Congenital VZV is associated with a high neonatal mortality rate (30%). Primary VZV infection during the first 6 months of pregnancy is associated with a 25% risk of in-utero infection (51). Twelve percent of intrauterine infections will result in a range of birth defects including limb hypoplasia, microcephaly, hydrocephaly, mental retardation and cataracts (51), in many ways similar to the disease spectrum currently observed with Zika fetal syndrome.

## Zika virus, a new viral TORCH pathogen

The list of TORCH viral pathogens is constantly expanding, and sufficient clinical data support adding ZIKV to the list. The exposure of a naïve population to a new virus which has historically been mosquito vectored, is sexually transmissible, and may be capable of direct human to human transmission by other means presents a greater challenge. With the emerging global threat of ZIKV infection to pregnant women, it is critical that we improve our understanding of the mechanism(s) of intrauterine infection, and of the medical management of subsequent neurologic disease.

Examination of the classic TORCH pathogens reveals some common themes, which can inform research concerning ZIKV fetal neuropathogenesis: these agents either infect the placenta, or infect specific tissues in the fetus linked to pathology. In some cases, specific molecular mechanisms that exacerbate the resulting pathology have been identified. Further exploration of cell surface receptors and placental permeability may assist with development of interventional prophylactics and therapeutics for pregnant women.

## Zika Virus Infection of the Placenta and Fetal Brain

In order to successfully establish an infection in a target tissue, all viruses must go through the same basic steps: the virus must overcome local host defenses at the site of infection (both barrier and immunologic response), infect a cell that is both susceptible and permissive to producing infectious virions, and the infected cell must release sufficient numbers of infectious particles which are able to travel to the target tissue and again infect a susceptible cell. Analyzing what we know about ZIKV infection in terms of this model can shed light on the possible mechanisms by which ZIKV might cause fetal abnormalities after initial maternal infection.

There are many plausible alternative hypotheses for Zika virus-induced fetal neuropathogenesis (100). These alternatives generally fall into two categories; infection of fetal tissue by ZIKV, or transcytosis of other factors that are causative of Zika Fetal Syndrome. Infection of fetal tissue may involve transcytosis of ZIKV from mother across the placenta or infection of the placenta itself. Either option may lead to dissemination of the virus in the fetus and subsequent infection of the developing brain. Infection of the placenta and resulting inflammatory response may indirectly alter neural development. Transcytosis of (yet to be defined) antigen-specific immunoglobulins or other maternal molecules related to the development of ZIKV GBS may directly harm the fetal brain without requiring viral replication in nervous tissue (15, 101, 102). ZIKV transfer and infection of the developing fetal brain may occur directly as free virus, as viral/non-neutralizing antibody complexes, or via infected Hofbauer or other migratory cells. Activation of TLR-3 by ZIKV binding to nervous tissue cells may directly induce damage without requiring viral replication (21). Placental infection by ZIKV triggering induction and release of inflammatory response-associated molecules may be sufficient to indirectly damage the fetal CNS (103-105). These possible mechanisms are not mutually exclusive, and may operate at different stages of fetal development.

The placenta represents a major barrier to fetal infection. This organ has evolved pathways for regulating the transport of materials, metabolites, oxygen and electrolytes, and both innate and adaptive immunologic effectors (particularly maternal immunoglobulin) between the mother and fetus. Soluble factors, oxygen and cells can all be selectively exchanged. Despite the relatively common event of infection of a pregnant woman by different viruses, transplacental passage of virus and intrauterine fetal infections are rare. This high degree of selectivity is largely due to a specialized outer placental layer; the syncytiotrophoblast, a large multinuclear body formed by the fusion of multiple cells into a syncytium during the second trimester of fetal development (106). This fusion into a single giant cell avoids the problems of maintaining intercellular junctions, which are sufficiently tight to prevent the unregulated movement of large molecules (and pathogens). In order for a virus to reach the fetus after this event, ZIKV must either have a mechanism to bypass the syncytiotrophoblast barrier, or must directly infect the placenta itself as has been observed with various viral TORCH pathogens. One possible method for the passage of ZIKV across the placenta to the fetus is through the mechanism which facilitates unidirectional transmission of maternal antibodies to the amniotic fluid and developing embryo (107, 108). The neonatal Fc receptor (FcRn, or FCGRT) is proposed to be involved in the recognition of maternal IgG, and in uptake of these antibodies by the cells of the infant gut. In addition, neonatal Fc gamma receptor IIb2 molecules expressed in human villous endothelium (within the FCGR2B2 compartment) actively participate in endothelial transcytosis of maternal IgG (109, 110). RAB3D, a member of the RAS-related protein RAB family, appears to play a key role in regulating the activity of the FCGR2B2 organelle, and therefore may influence transport of either autoimmune-associated antibodies or antibody-coated ZIKV. Antibody mediated enhancement of infection has been reported for Dengue virus, a related flavivirus, as well as for ZIKV (14). For Dengue virus, antibodies raised against previous infection with a different serotype of virus may enhance subsequent infection in a dendritic cell-mediated fashion (111, 112). For ZIKV, in vitro studies have demonstrated enhancement of infectivity with serum from patients with serologic responses to Dengue virus (14). The high degree of cross-reactivity between antibodies elicited by co-circulating arborviruses present in Brazil and throughout the Caribbean may contribute to intrauterine ZIKV disease by facilitating infected dendritic cell transport or by direct transcytosis of non-neutralizing antibody-coated ZIKV virions (14).

Delivery of ZIKV by transcytosis of antibody bound virus does not appear to be compatible with the window of greatest vulnerability for Zika teratogenicity, the first trimester of pregnancy. The transport of maternal IgG across the placenta begins at week sixteen (113, 114); the levels of IgG in fetal circulation at gestational weeks 17-22 are relatively low (5-10% of maternal levels) and rise continually with levels reaching 50% at weeks 28-32, followed by an exponential increase in the final four weeks before delivery (115). A study of RNA levels of Fc receptors in the placenta confirms that transcytosis is likely to begin primarily in the second trimester (116). Functionally active placental FcRn expression has been detected at 20 weeks (117). By analogy, maternal autoimmune antibody which may be elicited by ZIKV epitope mimics (ergo, GBS-associated antibodies) (118) are also unlikely to cross the placenta prior to the sixteenth gestational week. Many mothers of microcephalic children were infected with ZIKV before the tenth gestational week, and are likely to have cleared the virus well before sixteen weeks (29).

The timing of ZIKV infection relative to neonatal outcome may illuminate the mechanism of fetal infection. A recent preliminary report describes neuropathological aspects of fetal development in a cohort of Zika infected women (23). Most strikingly, fetal ultrasonography revealed abnormalities in twelve of the forty-two women who experienced ZIKV infection during pregnancy, as compared to none of the sixteen cohort-matched fetuses in Zika-negative women. Although the size of the cohort studied in this reported in this study was still low, they span a period of initial ZIKV exposure running from eight weeks to thirty-five weeks of gestation. The observations of microcephaly and severe cerebral pathology appear most commonly when the mother was infected with ZIKV at twelve weeks or earlier. Infection of the mother during the second or third trimester was reported to result in intrauterine growth restriction or, in two cases, fetal death. This pattern of timing supports the hypothesis that first trimester infection results in direct transmission of the virus to the fetal brain with subsequent viral replication, whereas later infection may involve activation of placental inflammatory responses. ZIKV infection of human cerebral organoids acts (at least in part) via TLR-3 to elicit a direct neural cell depletion which is partially abrogated by TLR-3 inhibition. TLR-3 activation by ZIKV resulted in alterations in expression of multiple genes associated with neuronal development, implying a mechanistic connection to disrupted neurogenesis (21).

The overall retardation of growth observed after second and third trimester exposure to ZIKV suggests that the virus may be exerting an indirect teratogenic effect by infecting the placenta rather than other fetal tissues during this period. A separate case study has recently identified infectious virus in the placenta of a fetus and detected resulting ongoing maternal ZIKV viremia (12), and this may include placental Hofbauer cell infection and/or activation (105). This is in agreement with previously published work showing that the placenta can induce viral resistance in nearby cells (119). In contrast, a well-designed basic virology study has shown that placental cells from a full term pregnancy are resistant to ZIKV (120). However, no data currently exist concerning the susceptibility of early placental cells to ZIKV infection.

Another possible mode of fetal infection would be transmission of ZIKV-infected maternal cells across the placenta at any stage of pregnancy. If a motile cell (such as a dendritic or Hofbauer cell) was infected and then crossed the placenta or was able to transit maternal-placental blood vessels, it could carry virus to the fetus. A similar situation has been modeled in mice in which dendritic cells can carry intracellular pathogens across the placenta (121). There is some limited evidence for the presence of maternal cells in the lymph nodes of second trimester fetuses, but the mechanism by which this migration occurs is not well understood (122). Infected migratory maternal cells might also contribute to fetal neuropathology via proinflammatory cytokine release. Placental Hofbauer cells have been shown to be activated by TLR-3 and TLR-4 mediated pathways, and ZIKV has been shown to activate TLR-3 mediated responses in neuronal cells (21).

Teratogenicity and neuropathology associated with TORCH pathogen infection of the placenta is well documented (54), and ZIKV may also interfere with fetal development by this route (103). The pronounced elevation of a variety of inflammatory cytokines may trigger microglial activation with attendant damage to surrounding cells, including neurophils, but is usually associated with damage to a wide range of fetal organs and tissue (123). The disease spectrum associated with chorioamnionitis overlaps with many of the features of Zika fetal syndrome, and includes periventricular leukomalacia, intraventricular hemorrhage, cerebral palsy, and retinopathy of prematurity (124-128). While ZIKV may also elicit similar pathology by direct placental infection, the striking selectivity and consistency of central nervous system damage observed, combined with the unusually severe damage to developing brain and the presence of ZIKV sequences in amniotic fluid and brain tissue, suggests some contribution of direct ZIKV infection of fetal CNS in the majority of cases.

## Expression of ZIKV receptors in placental and central nervous system tissues

Early in embryonic development, direct infection of the placenta by ZIKV could provide a route of entry to fetal tissue. Productive infection of the trophoblast by the virus would allow newly produced virions to be passed inward to the fetus. A critical step to the productive infection of any target cell is the expression of the correct viral receptors on the cell surface.

Flaviviruses, such as Dengue Virus (DV), Japanese Encephalitis Virus (JEV) and West Nile Virus (WNV) are known to use cellular C-type lectin proteins as receptors (129). Expression of several members of this receptor family is high on cells of the myeloid lineage such as monocytes, macrophages and dendritic cells (130). Multiple studies provide evidence for the role of one specific lectin, dendritic-cell specific ICAM-3-grabbing nonintegrin (DC-SIGN), in the infection of flaviviruses (131-135). DC-SIGN is an essential host protein that is involved in pathogen capture and antigen presentation in dendritic cells. As a lectin, DC-SIGN recognizes carbohydrate structures on proteins. Any ZIKV transmitted to a human host after replication in the salivary gland of a mosquito vector will carry the glycosylation pattern produced in the cells of the insect host. When virus replicates in insect salivary glands, the glycosylation of the viral proteins involved in receptor binding will follow the pattern observed in insects (high-mannose glycans) and not the more complex pattern seen in mammalian glycoproteins (132, 136). Dendritic cells are capable of recognizing this difference and reacting to these non-host glycosylation patterns. This specificity and the presence of dendritic cells in the epidermis, and therefore in close proximity to the site of the mosquito bite, means that mosquito-vectored flaviviruses are likely to preferentially infect the dendritic cell as an initial target cell type. The probability of uptake and initial infection of host dendritic cells may be enhanced by the presence of pre-existing non-neutralizing antibody which binds ZIKV (14).

Although the initial stages of human ZIKV infection are not as extensively studied as infection with viruses such as Dengue, a study by Hamel *et al*. has identified multiple receptors involved in ZIKV entry to the target cell (137). This seminal work examined the involvement of known Dengue virus receptors in ZIKV infection. The results confirmed a role for DC-SIGN in mediating ZIKV entry, and also identified roles for two TAM receptor proteins, called Tyro3 and AXL, and a minor role for a protein called TIM-1. Tyro3 and AXL are tyrosine kinase receptors whose natural ligand are the vitamin-K dependent proteins growth-arrest specific gene 6 (Gas6) and Protein S. Armed with this list of receptors, it is possible to predict what specific cells in the placenta and CNS might be susceptible to ZIKV infection.

An analysis from the US Centers for Disease Control and Prevention (CDC) reported ZIKV RNA and proteins in tissues from newborns and from two miscarriages (138). Examination of the corresponding placentas showed pathology associated with viral infection. Direct ZIKV infection of the placenta is plausible, as the trophoblast layer has been shown to express the needed receptors, and a recent report has recovered infectious virus from the placenta (12). AXL expression has been detected in the trophoblast, and perturbations in Gas6 signaling through AXL have been shown to be associated with pre-eclampsia, suggesting a possible mechanism of pathology (139). Histology available through the Human Protein Atlas also confirms expression of AXL and Tyro3 throughout the trophoblast layer (140). Although the trophoblast does not appear to express DC-SIGN, tissue resident cells of the myeloid lineage will express this lectin. This provides a pathway by which the infected trophoblast might produce virus that will infect patrolling myeloid cells. Infected myeloid cells may allow production of greater quantities of virus (leading to viremia) or serve as a vector to traffic virus to other tissues. Proof of this second possibility requires the identification of ZIKV positive perivascular macrophages or microglia in brain tissue from abortus specimens.

In order to selectively induce microcephaly and other observed changes in the brain, ZIKV must either alter pathways that affect CNS development or directly infect cells of the CNS. Comparisons to other viral TORCH pathogens strongly support the second possibility. It is worth noting that the early preparation of ZIKV in the laboratory setting was performed by intracerebral passage of the virus in neonatal mice. One study from 1971 presents an excellent microscopic examination of the brains of these mice (141). The authors catalog disruption of the pyriform cell layer of the Ammon’s horn and increased number of astrocytes without the presentation of infiltrating leukocytes. Examination of the tissue by electron microscopy reveals infected astroglia and neurons, but not microglia. The first indication that this was happening in humans involved histologic and molecular examination of products of conception including fetal brain tissue, which revealed the presence of viral particles in the brain of a fetus at 32 weeks of gestation (13). These findings have been supported and confirmed by a second paper examining another infected fetus (12). These case reports not only support the conclusion that the virus can replicate in cells of the CNS, but that the CNS serves as a site of viral persistence long after the mother was exposed. Again, the propensity for first trimester exposures to ZIKV provides clues about the possible mechanisms of neuropathogenesis. During the first trimester, the fetal blood brain barrier is ‘leaky’ and does not serve as a complete barrier against pathogens. Infection of the placenta in the first trimester and induction of fetal viremia may sufficiently disseminate virus, thereby enabling ZIKV access to the brain. Fetal development of a well-formed blood brain barrier later in pregnancy may also reduce the risk of CNS infection. A second possibility is that the frequency of target cells in the brain changes over time. A seminal report by Tang *et al*. reveals that ZIKV can infect neural progenitors (19) and this has been more recently confirmed in a study of ZIKV infection of human cerebral organoids in culture (18, 21). Infection of the brain in the first trimester might lead to infection of these precursor cells and associated pathology due to the ability of ZIKV to slow cellular replication and induce cell death. Supporting this hypothesis, direct examination of tissue from at least one ZIKV-positive fetus indicates that mature neurons are relatively unperturbed, suggesting that the progenitors may be preferentially infected (12). However, the reports by Bell *et al*. discussed above, as well as recent studies involving a more natural route of infection (142), demonstrate that infection of more mature brain cells is possible (141). Examination of the literature reveals the presence of Tyro3, AXL, DC-SIGN and TIM-1 on multiple cells in the CNS, leading to the hypothesis that multiple cell types might be infected (Table 2).

**Table 2.** **Expression of ZIKV receptors in human brain and placental tissue**. NA = data not available

|  | DC-SIGN | AXL | Tyro3 | TIM-1 | Evidence of Infection | References |
| --- | --- | --- | --- | --- | --- | --- |
| **CNS** |  |  |  |  |  |  |
| **Vascular Endothelial** | - | + | - | NA | Productive infection in tissue culture | (143-146) |
| **Perivascular macrophages** | NA | + | + | NA |  | (147) |
| **Astroglia** | - | + | + | NA | EM in mice | (143-146, 148, 149) |
| **Microglia** | - | + | + | NA |  | (143-146, 150) |
| **Neurons** | - | + | + | NA | EM in mice | (143-145, 149) |
| **Neuronal Precursors** | NA | NA | NA | NA | Productive infection in tissue culture | (19) |
| **Placenta** |  |  |  |  |  |  |
| **Trophoblast** | - | + | + | NA | Pathology | (139, 143) |
| **Dendritic Cells** | + | + | + | NA |  | (147, 151, 152) |

## Permissiveness to viral infection and alteration in cellular pathways

Not all cells expressing the receptor for a given virus are capable of being productively infected. The presence or absence of specific factors in the cell influence whether the virus can successfully establish an infection and produce more virus. At this time, little is known about the intracellular factors which may influence ZIKV replication. It may be that not all cells that display the appropriate receptors are capable of supporting viral replication. Genome wide RNAi screens have identified hundreds of cellular factors involved in flavivirus replication (153). Many of these factors are involved in critical host cell pathways such as: nucleic acid production, protein production and transport, lipid metabolism and energy production (153-155). Various interferon responsive genes have been shown to block flavivirus replication, as highlighted by the numerous mechanisms employed by the virus to counter these effects. However, in the absence of interferon, it is unclear if any cells are truly non-permissive to ZIKV infection.

What is clear is that flaviviruses have evolved multiple strategies for altering normal host cellular pathways to favor viral replication. Stress granules and P-bodies are accumulations of RNA found in the cytoplasm of cells that are involved in stress response, heat shock and response to infection by viruses (156, 157). Flaviviruses alter both of these granule types to increase viral replication. Interaction of viral non-coding regions with stress granule proteins has been implicated in increased viral RNA synthesis and processing of viral RNA by enzymes in the P-body, which leads to the accumulation of a non-coding viral RNA that may be involved in protecting the viral RNA against RNA interference (158, 159).

The existence of flavivirus encoded non-coding RNA (ncRNA) is of potential relevance to development of fetal neuropathology. The genome of ZIKV and other flaviviruses is relatively small. As such, there is evolutionary pressure to make efficient use of all available sequence to support viral replication and evasion of adaptive and innate host defenses. That the virus supports and maintains RNA and RNA structural motifs that are not directly used in the coding of proteins suggests that this non-coding RNA serves an important role in the viral life cycle (160). The production of ncRNA in flaviviruses is due to the incomplete digestion of viral RNA by XRN1, an exonuclease found in the P-body (159). Secondary structure in a stem loop within the untranslated region (UTR) prevent digestion of this area and leads to accumulation of viral ncRNA. Interestingly, this ncRNA seems to be essential for cytopathicity and viral pathogenesis. Viruses with mutations in the 3’UTR have no deficit in their ability to make viral RNA, but show attenuated cytopathic effects in infected cells. Two possible explanations have been given for this observation. The first is that the ncRNA modulates the host innate sensing proteins (Toll like receptors including TLR3, RIG-I and MDA5). Other studies show evidence that this ncRNA can function to inhibit the RNA interference pathway and alter the expression of host genes (161). When primary human fibroblasts are infected with Dengue virus, innate immune response signaling pathways are activated through both TLR3 and RIG- 1, but not Mda5, triggering up-regulation of IFNβ, TNFα, defensin 5 (HB5) and β defensin 2 (HβD2) (162). Heritable mutations in RIG-I and MDA5 coding sequences have been identified as causative for Type 1 interferonopathies (inherited autoimmune disorders associated with an inborn elevated interferon response) including Aicardi-Goutières syndrome, Systemic Lupus Erythematosus (SLE) in certain individuals as well as classic and atypical Singleton-Merten syndrome (163). As reviewed above, the radiographic characteristics of these syndromes overlap considerably with findings associated with both intrauterine CMV infection and Zika fetal syndrome. Prior assessment of therapeutic strategies for Aicardi-Goutières syndrome may help inform treatment options for Zika fetal syndrome (164). Hydroxychloroquine, used to treat SLE cerebritis and considered safe in pregnancy, is a potent inhibitor of Type I IFNs, and this therapeutic strategy may figure into the selection of drug-like entities being contemplated for treating pregnant women suffering from acute ZIKV (165-167).

Interactions of cellular proteins with the untranslated regions of the full length ZIKV RNA may also be critical for function. Examination of the West Nile Virus has shown that two cellular RNA-binding proteins, TIA-1 and TIAR, interact with the 3’ untranslated region (3’UTR) of that virus (158, 168). These proteins are essential host factors involved in formation of stress granules, and are sequestered at the site of viral RNA synthesis; an event that inhibits stress granule formation (168, 169). Viruses deficient in TIA-1 and TIAR binding replicate at a diminished rate in fibroblasts. A similar mechanism has been described for Dengue Virus (168). Due to the similarities to the secondary structure of the 3’UTR of these flaviviruses, ZIKV is likely to have similar effects. Whether ZIKV genomic or subgenomic RNA has binding sites for other host factors remains to be seen. Engagement of RNA-binding proteins specific to the brain or placenta by ZIKV might explain the pathology seen in the current epidemic.

The ability of ZIKV non-coding RNA to recruit cellular proteins might provide some insight into possible mechanisms of neuropathogenesis. The unique sequence of ZIKV may provide new targets for interaction with cellular proteins that are not seen in related viruses such as Dengue. Of particular interest will be whether factors specific to either the CNS or placenta bind to and regulate ZIKV RNA translation or replication. For example, the RNA binding protein Musashi-1 is expressed at high levels in neural precursors cells and can be found in both decidual and trophoblast cells in the placenta (140, 170).

Musashi-1 is required for differentiation and division of neural precursors, and is often used as a marker in identification of these cells (171, 172). Studies have revealed a role for Musashi as a regulator of mRNA translation, and that the protein is capable of both inhibiting and activating translation (173). Specifically, Musashi proteins play a role in regulating progenitor (stem) cell growth and differentiation through post-transcriptional control of gene expression (174). Musashi is also expressed in, and has been shown to influence mRNA translation in, a variety of epithelial stem cell types associated with glandular epithelium (174-177), spermatogenesis (178), brain and retinal tissue development (179, 180). Utilizing sequence alignment methods and available genomes of both historic and current ZIKV isolates, we have discovered a putative Musashi Binding Element (MBE) in the SL2 stem-loop of the 3’UTR (Figure 2) (181-184). Examination of ZIKV epidemic strains has revealed conserved changes in the NS2B open reading frame and 3’UTR relative to ancestral strains found in Africa (184). Our alignment confirms this, and highlights that two of these changes lie immediately upstream from the putative MBE. Both insects and mammals have Musashi homologs, and it has been reported that they bind MBE with slightly different sequence requirements (185). Application of the binding energy predictions of this work suggests that the evolutionary nucleotide polymorphism alterations observed in the region immediately upstream to the ZIKV core MBE may alter binding in mammals, but not the mosquito host. Given the expression of Musashi in neuronal precursors and the placenta, it will be critical to determine whether this element is involved in ZIKV pathogenesis, and if so, what ZIKV nucleotide polymorphisms may be associated with alterations in ZIKV Musashi Binding Element activity.

**Figure 2.**
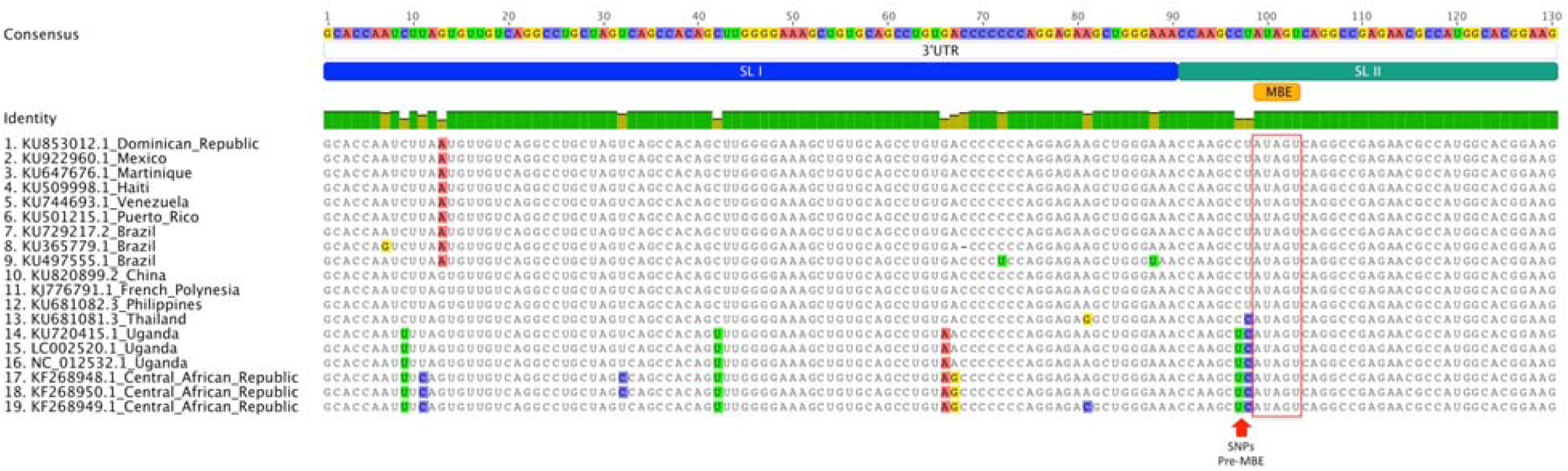
**Alignment of first 130 nucleotides of 3’UTR of ZIKV illustrating Musashi Binding Element location and associated mutations over time and geographic spread**. Sequences shown are the only that are unique for country and/or sequence, duplicates of same country were discarded. Alignment performed using MAFFT. Visualization using Geneious. Presence of SL I and SL II on those sequences, being SL II partially shown. Presence of Musashi Binding Element (MBE) on SL II, with two SNPs on African sequences, which could potentially change the RNA structure and availability of the element. SL I and SL II were annotated from Zhu Z. *et al*. MBE was annotated using the UTRscan tool of the UTRSite.

Flavivirus proteins insert themselves into the membrane of the endoplasmic reticulum (ER), forming invaginations that contain all of the proteins and RNA needed to produce additional viral RNA (186). These invaginations are connected to the cytoplasm by a small pore, through which the RNA is presumably passed to engage nearby ribosomes (187). Viral capsids are then assembled and enveloped by budding into the membranes of the Golgi. This dependence on membranes and the need to produce enough phospholipids to envelope all of the progeny virions has lead flaviviruses to evolve mechanism to alter membrane synthesis, lipid metabolism and ER processing (188-193).

The classic sign of flavivirus infection is the visualization by electron microscopy of small ‘viral factories’ where viral RNA and protein is made and then assembled into complete virions for release through the cellular transport system. It has been noted that these assemblages look very much like the autophagosomes formed during the process of autophagy. Autophagy is a normal cellular process wherein the cell digests large protein complex or intracellular pathogens, and has been shown to play an important role in the maintenance of stem cells (194). This process can provide a way for a cell to recycle materials under conditions of starvation or as a way to respond to intracellular infection (195). Studies of cells infected by ZIKV and other flaviviruses have shown an increase in the levels of autophagy (137, 196-198). Microscopic examination of intracellular compartments has revealed the presence of viral envelope protein (E protein) in the same vesicles as the autophagy marker LC3 (137). This suggests that the vesicles into which the virus buds may be autophagosomes. Some viruses block the late stages of autophagy, leading to the accumulation of autophagosomes that do not fuse with the lysosome. However, it seems that ZIKV does not block this step, and LC3 and E protein can be detected in mature autolysosomes. As the proper maturation of the viral envelope prior to release is pH dependent, it is possible that the virus has co-opted this pathway to maintain the correct pH and access proteases needed for maturation of the viral E protein. The trophoblast layer of the placenta produces miRNA that are pro-autophagic in nature, and which are delivered to bystander cells by exosomes (119). It is thought that this is a mechanism to make the trophoblast (and the cells in contact with it) more resistant to viral infection. However, in the case of ZIKV, this mechanism may help replication and spread by the virus once initial infection has been established, and could increase the susceptibility of nearby myeloid cells. Multiple lines of research suggest a role for autophagy in neurodegenerative diseases, which suggests that these ZIKV mediated changes in autophagy may also be involved in the observed neuropathic effects (195, 199, 200). Pharmacologic inhibition of autophagy is associated with inhibition of ZIKV replication in a variety of cell types, including human astrocytes (4, 137).

## Unanswered questions

In order to more completely understand the link between ZIKV infection and fetal abnormalities, more work must be done. The characteristic presentation of Zika Fetal Syndrome ranges from viral centric (microcephaly, blindness, ventricular calcifications and fetal presence of ZIKV by rt-PCR) to another extreme (long bone dysgenesis, negative for ZIKV) possibly associated with placental insufficiency. Epidemiological assessment of potential confounding risk factors for Zika fetal syndrome, including preceding immunologically cross-reactive arboviral infection and potential thalidomide sharing by patients being treated for leprosy, remains to be completed (14, 201, 202). To underscore the point, leprosy is now endemic throughout much of Brazil including Pernambuco (203), and post exposure prophylaxis of exposed individuals has been advocated (204, 205).

The gaps in understanding of ZIKV neuropathology highlighted in this review suggest that efforts should first be focused on obtaining clear, statistically significant data addressing a few specific questions. Prospective case control study reports on ZIKV infection of pregnant women and fetal outcomes are a step in the right direction. As such studies continue, a more definitive correlation between ZIKV infection and various congenital outcomes will become possible. Additionally, fundamental research will be required to answer questions regarding the ability of ZIKV to cross the placenta and infect the developing brain. Based on the published report of receptors utilized by ZIKV, a more complete survey of expression levels of these proteins in cells of the placenta should be prioritized. There is a desperate need for high quality histology and EM analysis of brain and placental tissue from different times after exposure. Although the Mlakar *et al*. report showed convincing evidence for the presence of viral particles in the brain of a thirty-two-week fetus, the method of fixation unfortunately makes it impossible to tell what specific cells may have been infected (13). A more recent analysis provides better clarity, but more studies will be needed (12). Some conclusions may be inferred from the work of Bell *et al*., but the injection of virus directly into the brain of neonatal mice may not be physiologically relevant (141). Recent progress involving the development and characterization of ZIKV infection using the AG129 mouse model are consistent with the findings of Bell *et al*., and may eventually enable a more complete understanding of the neural and glial tropism underlying ZIKV neuropathology (142). Although current literature provides some characterization of placental abnormalities, no definitive evidence has been shown supporting infection of specific cells of the placenta. A qualified animal model likely will be required to obtain this data. Finally, although PCR and histology are potentially powerful techniques, definitive proof of infection of a given tissue, or the relevance of virus reported in a biological sample, can only be obtained when replication competent virus can be retrieved from these samples.

To begin to understand the mechanism of ZIKV neuropathogenesis, other experiments might be considered. A survey of serum from ZIKV infected individuals could shed light on the development of self-reactive antibodies and possible links to GBS. Prior research and study designs which have illuminated the roles of viral proteins and regions or motifs of viral RNA in the pathogenesis of other flavivirus infections need to be applied to clarify the molecular virology of ZIKV. To what extent does ZIKV activation of TLR-3 contribute to fetal neuropathology? Are migratory placental cells such as Hofbauer cells infected by ZIKV during fetal development? Do specific proteins from the placenta and brain bind to the non-coding regions of ZIKV and play a role in the observed neural tissue disease? Recent studies have cataloged changes in the ZIKV genome as it has spread across the Pacific to the new world. Specific studies will be necessary to determine if these changes have in any way altered the transmissibility or virulence of the virus. Finally, the studied TORCH pathogens do not consistently cause pathology. It has been hypothesized that ZIKV infection may achieve access to the placenta and CNS secondary to some other event. Larger datasets will be needed to determine if ZIKV enters the fetus following some other perturbation, or whether other cofactors or confounding variables are associated with the severe congenital and adult neuropathology, which is now being observed with the current ZIKV outbreak in the Americas. But what is most clear is that ZIKV fetal neuropathology represents a new disease which does not completely overlap with the epidemiology or pathophysiology of other TORCH pathogens, and which will demand effort, resources, unparalleled collaboration, and above all, open mindedness in formulating public health responses as well as obstetrical and pediatric management strategies.

